## Supplementary material for "Divergent disulfide bond architecture defines two IgY subclasses in snakes": latex: main.tex

\documentclass{article}
\usepackage[utf8]{inputenc} % Recomendado para caracteres especiales
\usepackage[T1]{fontenc}
\usepackage{graphicx}
\usepackage{makecell}
\usepackage[numbers,sort&compress]{natbib} % sort&compress mejora la visualización [1-3]
\usepackage{float}
\usepackage{tikz}
\usetikzlibrary{arrows.meta,positioning,calc}
\usepackage{geometry} % Opcional: para mejores márgenes
\geometry{a4paper, margin=3.5cm}

% Ruta de las imágenes (ajusta si es necesario)
\graphicspath{ {./imagenes/} }

\title{Divergent disulfide bond architecture defines two IgY subclasses in snakes}
\author{Francisco Gambon Deza\\\small Independent researcher\\\small \texttt{}}
\date{January 2026}

\begin{document}

\maketitle

\begin{abstract}
Immunoglobulin Y (IgY) represents the major serum antibody in reptiles and birds, serving as the evolutionary precursor to mammalian IgG and IgE. While IgY diversification has been documented in several reptilian lineages, the structural basis underlying subclass divergence remains poorly understood. Here, we present a comprehensive phylogenetic and structural analysis of IgY sequences from 20 snake species, revealing two distinct evolutionary lineages (A and B) that arose through gene duplication. Structural modeling of the constant regions from \textit{Arizona elegans} identified a fundamental difference in the light chain-heavy chain (CL-CH1) disulfide bond architecture between lineages. Lineage B utilizes CYS16 in the CH1 domain (alignment position 13) for the inter-chain disulfide bond with the light chain CYS98, whereas Lineage A employs CYS136 (alignment position 99), representing N-terminal versus C-terminal positioning within the CH1 domain. Analysis of 50 diagnostic amino acid positions between lineages revealed that changes are distributed across all constant domains (CH1-CH4), with 13 positions showing radical substitutions affecting charge or polarity. Sliding window dN/dS analysis demonstrated purifying selection ($\omega < 1$) across both lineages, consistent with functional constraint following duplication. These findings provide structural evidence for subfunctionalization of snake IgY genes and suggest that alternative disulfide bond configurations may confer distinct biophysical or functional properties to each antibody subclass. This work advances our understanding of immunoglobulin evolution in reptiles and highlights the structural plasticity of antibody architecture.
\end{abstract}

\textbf{Keywords:} Immunoglobulin Y, IgY, snake immunology, gene duplication, subfunctionalization, disulfide bond, CL-CH1 interface, molecular evolution, Serpentes

\section{Introduction}

Immunoglobulin Y (IgY) is the predominant serum antibody class in birds, reptiles, and amphibians, functionally analogous to mammalian IgG \citep{warr1995igy}. Unlike the four-domain structure of mammalian IgG heavy chains (VH-CH1-CH2-CH3), IgY possesses an additional constant domain, resulting in a VH-CH1-CH2-CH3-CH4 architecture \citep{olivieri2021insights}. This structural difference, along with the absence of a flexible hinge region, confers distinct functional properties to IgY, including different effector functions and serum half-life compared to mammalian antibodies \citep{zhang2017igy}.

While IgY has been extensively characterized in birds, particularly in chickens (\textit{Gallus gallus}) due to its biotechnological applications \citep{karlsson2004chicken}, the immunoglobulin repertoire of reptiles remains poorly understood. Snakes (Serpentes) represent a particularly interesting group for immunological studies, as they have undergone significant adaptive radiation and occupy diverse ecological niches, from terrestrial to fully aquatic environments \citep{zheng2016combining}. Previous studies have identified IgY-like molecules in several snake species \citep{olivieri2021insights}, but a comprehensive analysis of their structural diversity and evolutionary history has been lacking.

Gene duplication is a major source of evolutionary innovation, allowing duplicated genes to acquire new functions (neofunctionalization) or partition ancestral functions (subfunctionalization) \citep{lynch2000evolutionary}. In the immunoglobulin superfamily, gene duplication has played a crucial role in generating the diversity of antibody classes and subclasses observed across vertebrates \citep{flajnik2010origin}. Understanding the evolutionary dynamics of duplicated immunoglobulin genes can provide insights into the functional diversification of the immune system.

The disulfide bond between the light chain constant region (CL) and the first heavy chain constant domain (CH1) is a fundamental structural feature of immunoglobulins, stabilizing the heterodimeric association between light and heavy chains \citep{nezlin1998immunoglobulins}. This inter-chain disulfide bond is highly conserved across vertebrate immunoglobulins, with the participating cysteine residues typically located at equivalent positions in the primary sequence. Alterations in this structural feature could have significant implications for antibody assembly, stability, and function.

In this study, we performed a comprehensive phylogenetic and structural analysis of IgY heavy chain constant regions from 20 snake species representing major families including Colubridae, Viperidae, Elapidae, Pythonidae, and Boidae. Our analysis reveals the existence of two distinct evolutionary lineages of snake IgY that differ fundamentally in their light chain-heavy chain disulfide bond architecture. Furthermore, we investigated the selective pressures acting on these lineages using dN/dS analysis, providing insights into the evolutionary forces that have shaped and maintained this structural divergence.

Our findings demonstrate that:
\begin{enumerate}
    \item Snake IgY genes have undergone an ancient duplication event, giving rise to two distinct lineages (Lineage A and Lineage B) that predate the radiation of advanced snakes (Colubroidea).
    \item The two lineages utilize different cysteine residues for the CL-CH1 disulfide bond: position 99 (N-terminus of CH1) in Lineage A versus position 13 (C-terminus of CH1) in Lineage B.
    \item The CH1 domain, particularly the regions containing the diagnostic cysteines, is under strong purifying selection, confirming the functional importance of the disulfide bond architecture.
    \item The CH3 domain shows strong signals of positive selection ($\omega$ up to 4.5), suggesting active functional diversification between the lineages.
    \item The divergence between lineages has been driven by positive selection ($\omega$ = 1.57), consistent with subfunctionalization following gene duplication.
\end{enumerate}

These results provide the first evidence of alternative disulfide bond architectures in vertebrate immunoglobulins and suggest that the two snake IgY lineages may have acquired distinct functional properties during their evolutionary divergence.

\section{Materials and Methods}

\subsection{Sequence Data Collection}

IgY heavy chain constant region sequences were obtained from 20 snake species representing diverse families within Serpentes. A total of 40 sequences were compiled, including both previously published sequences and novel annotations from genomic databases. Sequences encompassed the complete constant region domains (CH1-CH2-CH3-CH4) without the variable region (VH). Species representation included members from Colubridae, Viperidae, Elapidae, and Boidae families, ensuring broad taxonomic coverage across snake phylogeny.

\subsection{Sequence Alignment and Phylogenetic Analysis}

Multiple sequence alignment of amino acid sequences was performed using MAFFT v7.505 \citep{katoh2013mafft} with default parameters. The alignment was filtered to retain positions with $\geq$80\% occupancy, resulting in 417 aligned positions. Phylogenetic reconstruction was conducted using the neighbor-joining method implemented in BioPython \citep{cock2009biopython}, with bootstrap support calculated from 1,000 replicates. The resulting tree was visualized and annotated using custom Python scripts with the ETE3 toolkit \citep{huerta2016ete3}.

\subsection{Codon Alignment and Selection Analysis}

Nucleotide sequences corresponding to the filtered amino acid alignment were extracted and codon-aligned using PAL2NAL \citep{suyama2006pal2nal} to maintain reading frame integrity. The codon alignment comprised 417 codons (1,251 nucleotides) across all 40 sequences, with 0\% translation errors verified against the amino acid alignment.

Evolutionary selection pressures were assessed using sliding window analysis of nonsynonymous to synonymous substitution ratios (dN/dS, $\omega$). Windows of 30 codons with a step size of 5 codons were analyzed across the alignment, calculating pairwise dN/dS values using the Nei-Gojobori method \citep{nei1986simple}. Global $\omega$ values were computed for within-lineage and between-lineage comparisons to assess differential selection pressures.

\subsection{Structural Modeling and Analysis}

Three-dimensional structures of IgY constant regions were predicted using AlphaFold2 \citep{jumper2021highly} for representative sequences from \textit{Arizona elegans}. Structures were modeled as heterotetramers comprising two light chain constant domains (CL) and two heavy chain constant regions (CH1-CH2-CH3-CH4), reflecting the native antibody architecture. Model quality was assessed using predicted Local Distance Difference Test (pLDDT) scores.

Structural analysis and visualization were performed using Py3Dmol \citep{rego2015py3dmol} for interactive 3D representations and matplotlib \citep{hunter2007matplotlib} for publication-quality figures. Disulfide bond identification was conducted by measuring inter-residue distances between cysteine sulfur atoms, with bonds assigned when S$\gamma$-S$\gamma$ distances were $<$3.0 \AA.

\subsection{Diagnostic Position Identification}

Lineage-specific diagnostic positions were identified by comparing amino acid frequencies between Lineage A and Lineage B sequences at each aligned position. A position was considered diagnostic if: (1) the consensus amino acid differed between lineages, and (2) the conservation frequency exceeded 70\% in at least one lineage. Amino acid changes were classified as conservative or radical based on physicochemical properties including charge, polarity, and side chain volume \citep{grantham1974amino}.

Diagnostic positions were mapped to the three-dimensional structures using local sequence alignment (Smith-Waterman algorithm) to account for length differences between the filtered alignment and structural coordinates. Positions were assigned to immunoglobulin domains (CH1, CH2, CH3, CH4) based on canonical domain boundaries.

\subsection{Disulfide Bond Interface Analysis}

The CL-CH1 interface was analyzed to identify cysteine residues involved in inter-chain disulfide bond formation. For each lineage structure, all cysteine pairs between the light chain (chains A/B) and heavy chain CH1 domain (chains C/D) were evaluated. Inter-chain disulfide bonds were assigned based on: (1) S$\gamma$-S$\gamma$ distance $<$3.0 \AA, (2) appropriate geometry for disulfide bond formation, and (3) surface accessibility at the CL-CH1 interface.

\subsection{Data Visualization}

Phylogenetic trees were visualized with lineage-specific coloring (Lineage A: blue; Lineage B: red). Three-dimensional molecular visualizations were generated as interactive HTML files using Py3Dmol, with diagnostic positions highlighted using sphere representations. Publication-quality figures were exported in both PNG (300 dpi) and SVG formats for editorial flexibility.

\subsection{Software and Data Availability}

All analyses were performed using Python 3.10 with the following packages: BioPython \citep{cock2009biopython}, NumPy \citep{harris2020array}, Pandas \citep{mckinney2010data}, Matplotlib \citep{hunter2007matplotlib}, and Py3Dmol \citep{rego2015py3dmol}. Sequence alignments, phylogenetic trees, and structural coordinates are available upon request.

In addition, Codex (v5.2) was used as an AI-assisted writing and editing tool to support drafting and language polishing of the manuscript text. All scientific content, analyses, and interpretations were defined and validated by the authors.

In addition, Codex (v5.2) was used as an AI-assisted writing and editing tool to support drafting and language polishing of the manuscript text. All scientific content, analyses, and interpretations were defined and validated by the author.

\section*{Results}

\subsection{Processed alignment and phylogenetic structure of snake IgY}

All downstream analyses were based on the processed snake IgY constant-region alignment (\texttt{IgY\_all\_aa.aln.occ80.fasta}), which contains translated sequences derived from the CH1--CH4 exons and was filtered to remove columns with $>$80\% gaps.

\begin{figure}[H]
\centering
\includegraphics[width=0.95\linewidth]{IgY_all_aa.aln.occ80.fasta.treefile.pdf}
\caption{Phylogenetic tree inferred from the filtered snake IgY constant-region alignment. The tree supports two IgY lineages in snakes: Lineage~A (blue) and Lineage~B (red).}
\label{fig:igy_tree}
\end{figure}

Phylogenetic reconstruction using this filtered alignment recovered two clearly separated IgY lineages in snakes (Figure~\ref{fig:igy_tree}). These correspond to a Lineage~A (highlighted in blue) and a Lineage~B (highlighted in red), consistent with a deep split within snake IgY constant regions. Notably, the genomic position of individual C$\upsilon$ (IgY) genes relative to IgD does not appear to be lineage-specific across species: in some loci the first IgY gene downstream of IgD (IgY1) falls within Lineage~A, whereas in others IgY1 clusters within Lineage~B, suggesting an apparently non-conserved (or reshuffled) arrangement of C$\upsilon$ genes with respect to lineage identity.

Phylogenetic analysis of 40 IgY heavy chain constant-region sequences (CH1--CH4) from 20 snake species revealed two distinct evolutionary lineages (Lineage~A: 23 sequences; Lineage~B: 17 sequences). Both lineages are present in Colubridae, Viperidae, and Elapidae, suggesting that the gene duplication event predates the radiation of advanced snakes (Colubroidea). Notably, the within-species nomenclature IgY\_1/IgY\_2 does not correspond to lineage identity across taxa, indicating that paralog numbering is not evolutionarily conserved.

\subsection{Structural divergence at the CL-CH1 interface}

\begin{figure}[H]
\centering
\includegraphics[width=0.9\linewidth]{IgY1.png}\\[0.6em]
\includegraphics[width=0.9\linewidth]{IgY2.png}
\caption{Representative constant-region structural models for the two snake IgY lineages. Top: Lineage~A model; bottom: Lineage~B model.}
\label{fig:igy12_stack}
\end{figure}

The distinct disulfide bond architecture between the two lineages implies different relative orientations of the light and heavy chains, potentially affecting molecular flexibility and function. This structural divergence, combined with the presence of both lineages across multiple snake families, suggests an ancient gene duplication followed by subfunctionalization, with each lineage acquiring distinct structural characteristics that have been maintained throughout colubroid evolution.

\begin{figure}[H]
    \centering
    \includegraphics[width=0.85\textwidth]{Figure_IgY_disulfide_scheme_publication.pdf}
    \caption{\textbf{Differential disulfide bond architecture between snake IgY lineages.} Schematic representation of the light chain constant domain (CL) and heavy chain CH1 domain showing the distinct cysteine positions involved in the inter-chain disulfide bond.}
    \label{fig:disulfide_scheme}
\end{figure}

Interestingly, pythons and boas (more basal snake lineages) appear to possess only Lineage A-type sequences, suggesting that Lineage B may represent a derived innovation specific to advanced snakes. The case of \textit{Charina bottae}, which possesses cysteines at both positions 13 and 99, may represent either an ancestral or intermediate state warranting further investigation.

\subsection{Evolutionary Selection Analysis}

To investigate the evolutionary forces shaping the divergence between the two IgY lineages, we performed dN/dS ($\omega$) analysis using a sliding window approach (30 codons, step size 5) across the heavy chain constant region alignment.

\subsubsection{Global dN/dS Analysis}

Pairwise dN/dS calculations revealed distinct selective pressures acting on each lineage (Table \ref{tab:dnds_global} and Figure \ref{fig:dnds_sliding}):

\begin{table}[h]
\centering
\caption{Global dN/dS values within and between IgY lineages}
\label{tab:dnds_global}
\begin{tabular}{lcccl}
\hline
Comparison & dN & dS & $\omega$ (dN/dS) & Interpretation \\
\hline
Within Lineage A & 0.161 & 0.156 & 1.13 & Weak positive selection \\
Within Lineage B & 0.207 & 0.228 & 0.92 & Approximately neutral \\
Between lineages & 0.499 & 0.320 & 1.57 & Strong positive selection \\
\hline
\end{tabular}
\end{table}

The elevated $\omega$ value between lineages (1.57) suggests that the divergence between Lineage A and Lineage B has been driven by positive selection, consistent with subfunctionalization following gene duplication.

\subsubsection{Domain-Specific Selection Patterns}

\begin{figure}[H]
    \centering
    \includegraphics[width=\textwidth]{dNdS_sliding_window.png}
    \caption{\textbf{Selection pressure along the IgY heavy chain constant region.} Sliding window analysis (30 codons, step size 5) of dN/dS ($\omega$) across the CH1-CH4 domains.}
    \label{fig:dnds_sliding}
\end{figure}

Sliding window analysis revealed heterogeneous selection pressures across the constant region (Table \ref{tab:dnds_domain}, Figure \ref{fig:dnds_sliding}):

\begin{table}[h]
\centering
\caption{Domain-specific dN/dS values in snake IgY}
\label{tab:dnds_domain}
\begin{tabular}{lccl}
\hline
Domain & Mean $\omega$ & Range & Interpretation \\
\hline
CH1 & 0.82 & 0.36--1.51 & Purifying selection (conserved) \\
CH2 & 1.20 & 0.25--2.89 & Mixed selection \\
CH3 & 2.18 & 0.80--4.46 & Strong positive selection \\
CH4 & 1.89 & 1.08--2.67 & Moderate positive selection \\
\hline
\end{tabular}
\end{table}

Notably, CH3 exhibited the strongest signal of positive selection ($\omega$ up to 4.46), suggesting active functional diversification in this domain. This region may represent a site of interaction with lineage-specific receptors or ligands.

\subsubsection{Selection at Diagnostic Cysteine Positions}

The diagnostic cysteine positions involved in the CL-CH1 disulfide bond showed distinct selection patterns:

\begin{itemize}
    \item Position 13 (Cys in Lineage B): Located within a region of purifying selection ($\omega$ = 0.35--0.47)
    \item Position 99 (Cys in Lineage A): Located within a region of purifying selection ($\omega$ = 0.46--0.82)
    \item Position 210 (Cys in Lineage B): Located at the CH2/CH3 boundary, transitioning to positive selection
\end{itemize}

The strong purifying selection acting on the regions containing the diagnostic cysteines confirms their functional importance in maintaining the structural integrity of the light chain-heavy chain interface. Despite the different positions used for the disulfide bond in each lineage, both are maintained under strong selective constraint.

\subsubsection{Conclusions}

The selection analysis supports a model of subfunctionalization following an ancient gene duplication event. While the CL-CH1 interface (CH1 domain) remains under purifying selection to maintain structural integrity, the CH3 and CH4 domains have undergone positive selection, potentially acquiring lineage-specific functions. The distinct disulfide bond architectures in each lineage, combined with the strong purifying selection at these sites, suggest that this structural difference was established early in the divergence of the two lineages and has been maintained by natural selection.

\section{Discussion}

\subsection{Evidence for IgY Gene Duplication in Snakes}

Our phylogenetic analysis of 40 IgY sequences from 20 snake species reveals a clear bifurcation into two distinct evolutionary lineages, designated Lineage A and Lineage B. This pattern is consistent with an ancient gene duplication event that occurred early in snake evolution, followed by lineage-specific divergence. The presence of both lineages across diverse snake families (Colubridae, Viperidae, Elapidae, and Boidae) suggests that this duplication predates the radiation of modern snake families, indicating a duplication event at least 60-80 million years ago \citep{olivieri2021insights}.

Gene duplication is a major source of evolutionary novelty, providing raw genetic material for functional diversification \citep{ohno1970evolution, lynch2000evolutionary}. Following duplication, paralogous genes may undergo several fates: nonfunctionalization (pseudogenization), neofunctionalization (acquisition of new functions), or subfunctionalization (partitioning of ancestral functions) \citep{force1999preservation, birchler2012gene}. The retention of both IgY lineages across snake phylogeny, combined with evidence of purifying selection in both lineages, strongly supports a subfunctionalization model rather than neutral drift or neofunctionalization.

\subsection{Structural Divergence at the CL-CH1 Interface}

The most striking finding of this study is the identification of fundamentally different disulfide bond architectures between the two IgY lineages. In Lineage B, the inter-chain disulfide bond connecting the light chain constant domain (CL) to the heavy chain CH1 domain utilizes CYS16 of CH1 (alignment position 13), located in the N-terminal region of the domain. In contrast, Lineage A employs CYS132 (alignment position 99), positioned in the C-terminal region of CH1. This represents a spatial displacement of approximately 86 amino acid positions along the CH1 domain.

The CL-CH1 disulfide bond is critical for antibody stability and function, covalently linking the light and heavy chains to form the functional Fab region \citep{liu2012disulfide, schroeder2010structure}. In mammalian immunoglobulins, the position of this bond is highly conserved, with IgG subclasses showing consistent CL-CH1 connectivity \citep{wypych2008human}. The discovery of alternative disulfide bond positions in snake IgY represents, to our knowledge, the first documented case of such structural plasticity in the CL-CH1 interface among vertebrate immunoglobulins.

This structural divergence may have functional implications. The N-terminal versus C-terminal positioning of the inter-chain bond could affect: (1) the relative orientation of VL and VH domains, potentially influencing antigen binding geometry; (2) the flexibility of the Fab arm, affecting avidity and cross-linking capacity; and (3) the stability of the CL-CH1 interface under different physiological conditions \citep{stanfield2006antibody, sela2013structural}.

\subsection{Evolutionary Implications of Alternative Disulfide Architectures}

The existence of two distinct disulfide bond configurations raises intriguing questions about the evolutionary trajectory of this structural feature. Two scenarios merit consideration:

First, the ancestral snake IgY may have possessed both cysteine positions, with each lineage subsequently losing one through mutation. This would represent a case of complementary degenerative mutations leading to subfunctionalization, as proposed by the DDC (Duplication-Degeneration-Complementation) model \citep{force1999preservation}. Under this scenario, the ancestral gene may have had redundant structural features that were partitioned between duplicates.

Alternatively, one disulfide configuration may represent the ancestral state, with the other arising through neofunctionalization in one lineage. Comparison with IgY sequences from other reptilian groups (crocodilians, turtles, lizards) and birds could help resolve which configuration is ancestral \citep{zhang2017igy, xu2009igm}.

\subsection{Functional Diversification and Immune Repertoire}

The maintenance of two structurally distinct IgY subclasses in snakes likely reflects functional diversification that enhances immune capabilities. Similar subclass diversification has been observed in mammalian IgG, where different subclasses exhibit distinct effector functions, half-lives, and tissue distributions \citep{vidarsson2014igg}. In birds, IgY truncation variants (IgY$\Delta$Fc) have been described that lack the Fc region and may serve specialized functions \citep{warr1995igy, zhang2017igy}.

The structural differences between snake IgY lineages could translate into functional differences in several ways. Different CL-CH1 geometries might affect: (1) complement activation efficiency; (2) Fc receptor binding affinity; (3) serum half-life through differential FcRn interactions; or (4) mucosal versus systemic distribution. Experimental characterization of recombinant IgY from both lineages would be required to test these hypotheses.

\subsection{Selection Pressures and Functional Constraint}

Our sliding window dN/dS analysis revealed that both IgY lineages are under purifying selection ($\omega < 1$) across the constant region domains. This pattern indicates that both subclasses are functionally constrained and not evolving neutrally, consistent with their retention serving adaptive purposes. The absence of strong positive selection signals suggests that the major functional divergence occurred early after duplication, with subsequent evolution focused on maintaining established functions rather than acquiring new ones.

Interestingly, the dN/dS values showed some variation across domains, with certain regions exhibiting stronger purifying selection than others. The CH1 domain, which harbors the divergent disulfide bond positions, showed intermediate selection pressure, possibly reflecting a balance between structural constraint and adaptive flexibility at the CL-CH1 interface.

\subsection{Comparative Context: Immunoglobulin Evolution in Reptiles}

Our findings contribute to the growing understanding of immunoglobulin diversity in reptiles, a group that has been historically understudied compared to mammals and birds \citep{zimmerman2010reptilian, merchant2013immunology}. Previous work has documented IgY subclass diversity in snakes \citep{gambon2012snakes} and identified novel immunoglobulin isotypes in lizards and crocodilians \citep{magadan2013igh}. The present study extends these observations by providing structural evidence for the molecular basis of subclass divergence.

Reptiles occupy a key phylogenetic position for understanding immunoglobulin evolution, representing the transition between the IgY-dominated systems of fish, amphibians, and birds, and the IgG/IgE/IgA diversification seen in mammals \citep{flajnik2010origin}. The structural plasticity observed in snake IgY suggests that the immunoglobulin fold can accommodate significant variation in inter-chain connectivity while maintaining overall function, a finding with implications for understanding antibody evolution more broadly.

\subsection{Limitations and Future Directions}

Several limitations of this study should be acknowledged. First, the three-dimensional structures analyzed were computational predictions from AlphaFold2 rather than experimentally determined structures. While AlphaFold2 has demonstrated remarkable accuracy for immunoglobulin domains \citep{jumper2021highly}, experimental validation through X-ray crystallography or cryo-EM would strengthen our structural conclusions.

Second, our analysis focused on the constant regions and did not include variable region diversity, which is critical for antigen recognition. Future studies integrating VH and VL repertoire analysis with constant region subclass assignment would provide a more complete picture of snake antibody diversity.

Third, functional characterization of the two IgY lineages remains to be performed. Expression of recombinant IgY from both lineages, followed by biophysical characterization and functional assays (complement activation, Fc receptor binding, stability measurements), would directly test whether the structural differences translate into functional divergence.

\subsection{Conclusions}

This study provides structural evidence for subfunctionalization following IgY gene duplication in snakes. The identification of alternative disulfide bond architectures at the CL-CH1 interface represents a novel finding that expands our understanding of immunoglobulin structural plasticity. The conservation of both lineages across snake phylogeny, combined with signatures of purifying selection, indicates that both subclasses serve important, non-redundant functions in snake immunity. These findings highlight the value of studying immunoglobulin evolution in non-model organisms and suggest that significant structural diversity remains to be discovered in vertebrate antibody systems.

\bibliographystyle{model5} % Asegúrate de tener el archivo model5.bst
\bibliography{references}

\clearpage
\section*{Supplementary Information}
\addcontentsline{toc}{section}{Supplementary Information}

The amino-acid alignment used to guide all downstream analyses in this study is provided as Supplementary Information. The file, named \texttt{IgY\_all\_aa.aln.occ80.fasta}, is a plain-text FASTA-formatted multiple sequence alignment (MSA) of snake IgY constant-region sequences. It corresponds to the occupancy-filtered alignment (columns retained at $\geq$80\% occupancy) used for phylogenetic reconstruction and all subsequent comparative and selection analyses. The alignment can be opened and inspected with any standard text editor, or imported into common sequence-analysis software for visualization and downstream processing.


\end{document}
