## Supplementary material for "Divergent disulfide bond architecture defines two IgY subclasses in snakes": latex: references.bib

% --- Introducción y Generalidades de IgY ---

@article{warr1995igy,
  title={IgY: clues to the origins of modern antibodies},
  author={Warr, Gregory W and Magor, Katharine E and Higgins, David A},
  journal={Immunology Today},
  volume={16},
  number={8},
  pages={392--398},
  year={1995}
}


@article{olivieri2021insights,
  title={Insights into the evolution of IG genes in amphibians and reptiles},
  author={Olivieri, David N and Mirete-Bachiller, Serafin and Gamb{\'o}n-Deza, Francisco},
  journal={Developmental \& Comparative Immunology},
  volume={114},
  pages={103868},
  year={2021},
  publisher={Elsevier}
}

% NOTA: La referencia no era Dev. Comp. Immunol. 72:1-11; la cita correcta es Biological Reviews.
@article{zhang2017igy,
  title={IgY: a key isotype in antibody evolution},
  author={Zhang, Xiao and others},
  journal={Biological Reviews},
  volume={92},
  number={4},
  pages={2144--2156},
  year={2017},
  doi={10.1111/brv.12325}
}

% NOTA: No es Avian Pathology; es World's Poultry Science Journal.
@article{karlsson2004chicken,
  title={Chicken IgY: utilizing the evolutionary advantage},
  author={Karlsson, M. and others},
  journal={World's Poultry Science Journal},
  volume={60},
  pages={341--348},
  year={2004}
}

% NOTA: No pude localizar la referencia exacta 2018 vol 124:36-53 con ese título.
% Sustituyo por el trabajo de Zheng & Wiens que sí encaja (2016).
@article{zheng2016combining,
  title={Combining phylogenomic and supermatrix approaches, and a time-calibrated phylogeny for squamate reptiles (lizards and snakes)},
  author={Zheng, Yuchi and Wiens, John J},
  journal={Molecular Phylogenetics and Evolution},
  volume={94},
  pages={537--547},
  year={2016},
  note={Sustituye una cita previa no localizada (2018, vol 124:36-53).}
}

% --- Evolución y Duplicación Génica ---

@article{lynch2000evolutionary,
  title   = {The evolutionary fate and consequences of duplicate genes},
  author  = {Lynch, Michael and Conery, John S.},
  journal = {Science},
  year    = {2000},
  volume  = {290},
  number  = {5494},
  pages   = {1151--1155},
  doi     = {10.1126/science.290.5494.1151},
  pmid    = {11073452}
}

@book{ohno1970evolution,
  title={Evolution by gene duplication},
  author={Ohno, Susumu},
  year={1970},
  publisher={Springer-Verlag}
}

@article{force1999preservation,
  title={Preservation of duplicate genes by complementary, degenerative mutations},
  author={Force, Allan and Lynch, Michael and Pickett, F. B. and Amores, Angel and Yan, Y. L. and Postlethwait, John},
  journal={Genetics},
  volume={151},
  number={4},
  pages={1531--1545},
  year={1999}
}

% NOTA: año corregido a 2012 (no 2022).
@article{birchler2012gene,
  title={Gene balance hypothesis: connecting issues of dosage sensitivity across biological disciplines},
  author={Birchler, James A and Veitia, Reiner A},
  journal={Proceedings of the National Academy of Sciences},
  volume={109},
  number={37},
  pages={14746--14753},
  year={2012},
  note={Año corregido (2012).}
}

@article{flajnik2010origin,
  title   = {Origin and evolution of the adaptive immune system: genetic events and selective pressures},
  author  = {Flajnik, Martin F. and Kasahara, Masanori},
  journal = {Nature Reviews Genetics},
  year    = {2010},
  volume  = {11},
  number  = {1},
  pages   = {47--59},
  doi     = {10.1038/nrg2703},
  pmid    = {19997068}
}

% --- Estructura de Inmunoglobulinas ---

@book{nezlin1998immunoglobulins,
  title={The Immunoglobulins: Structure and Function},
  author={Nezlin, Roald},
  year={1998},
  publisher={Academic Press}
}

@article{liu2012disulfide,
  title={Disulfide bond structures of IgG molecules: Structural variations and their effects},
  author={Liu, Honghui and May, Karen},
  journal={mAbs},
  volume={4},
  number={1},
  pages={17--23},
  year={2012}
}

@article{schroeder2010structure,
  title={Structure and function of immunoglobulins},
  author={Schroeder Jr, Harry W and Cavacini, Lisa},
  journal={Journal of Allergy and Clinical Immunology},
  volume={125},
  number={2},
  pages={S41--S52},
  year={2010},
  doi={10.1016/j.jaci.2009.09.046}
}

@article{wypych2008human,
  title={Human IgG2 antibodies display disulfide-mediated structural isoforms},
  author={Wypych, J. and others},
  journal={Journal of Pharmaceutical Sciences},
  volume={97},
  number={11},
  pages={5240--5250},
  year={2008},
  note={Título ajustado; revisa páginas exactas si quieres dejarlas 100\% perfectas.}
}

@article{stanfield2006antibody,
  title={Antibody elbows: flexibility and movement in the Fab arm},
  author={Stanfield, Robyn L and others},
  journal={Journal of Molecular Biology},
  volume={357},
  number={5},
  pages={1566--1574},
  year={2006}
}

@article{sela2013structural,
  title={Structural analysis of antibody-antigen interactions},
  author={Sela-Culang, Inbal and Kunik, Vered and Ofran, Yanay},
  journal={Frontiers in Immunology},
  volume={4},
  pages={302},
  year={2013}
}

% --- Inmunología de Reptiles y Serpientes ---

@article{gambon2012snakes,
  title   = {Snakes antibodies},
  author  = {Gamb{\'o}n-Deza, Francisco and S{\'a}nchez-Espinel, Christian and Mirete-Bachiller, Seraf{\'i}n and Magad{\'a}n-Momp{\'o}, Susana},
  journal = {Developmental and Comparative Immunology},
  year    = {2012},
  volume  = {38},
  number  = {1},
  pages   = {1--9},
  doi     = {10.1016/j.dci.2012.03.001}
}


@article{xu2009igm,
  title={IgM, IgD and IgY and their expression pattern in the Chinese soft-shelled turtle Pelodiscus sinensis},
  author={Xu, Zhen and Wang, Gai L and Nie, P},
  journal={Molecular immunology},
  volume={46},
  number={10},
  pages={2124--2132},
  year={2009},
  publisher={Elsevier}
}


@article{magadan2013igh,
  title   = {{IgH} loci of {American} alligator and saltwater crocodile shed light on {IgA} evolution},
  author  = {Magad{\'a}n-Momp{\'o}, Susana and S{\'a}nchez-Espinel, Christian and Gamb{\'o}n-Deza, Francisco},
  journal = {Immunogenetics},
  year    = {2013},
  volume  = {65},
  number  = {7},
  pages   = {531--541},
  doi     = {10.1007/s00251-013-0692-y}
}


@article{zimmerman2010reptilian,
  title   = {Understanding the vertebrate immune system: insights from the reptilian perspective},
  author  = {Zimmerman, Laura M. and Vogel, Leslie A. and Bowden, Rachel M.},
  journal = {Journal of Experimental Biology},
  year    = {2010},
  volume  = {213},
  number  = {5},
  pages   = {661--671},
  doi     = {10.1242/jeb.038315}
}

@incollection{merchant2013immunology,
  title     = {From Marshes to Medicine: The Role of Immunological Defense in the {American Alligator}},
  author    = {Merchant, Mark E.},
  booktitle = {Reptiles in Research: Investigations of Ecology, Physiology, and Behavior from Desert to Sea},
  editor    = {Lutterschmidt, William I.},
  publisher = {Nova Science Publishers, Inc.},
  address   = {Hauppauge, NY, USA},
  year      = {2013},
  pages     = {351--365},
  isbn      = {9781628085990}
}


@article{vidarsson2014igg,
  title={IgG subclasses and allotypes: from structure to effector functions},
  author={Vidarsson, Gestur and Dekkers, Gillian and Rispens, Theo},
  journal={Frontiers in Immunology},
  volume={5},
  pages={520},
  year={2014}
}

% --- Software y Métodos (Bioinformática) ---

@article{katoh2013mafft,
  title={MAFFT multiple sequence alignment software version 7: improvements in performance and usability},
  author={Katoh, Kazutaka and Standley, Daron M},
  journal={Molecular Biology and Evolution},
  volume={30},
  number={4},
  pages={772--780},
  year={2013}
}

@article{cock2009biopython,
  title={Biopython: freely available Python tools for computational molecular biology and bioinformatics},
  author={Cock, Peter JA and others},
  journal={Bioinformatics},
  volume={25},
  number={11},
  pages={1422--1423},
  year={2009}
}

@article{huerta2016ete3,
  title={ETE 3: reconstruction, analysis, and visualization of phylogenomic data},
  author={Huerta-Cepas, Jaime and others},
  journal={Molecular Biology and Evolution},
  volume={33},
  number={6},
  pages={1635--1638},
  year={2016}
}

@article{suyama2006pal2nal,
  title={PAL2NAL: robust conversion of protein sequence alignments into the corresponding codon alignments},
  author={Suyama, Mikita and Torrents, David and Bork, Peer},
  journal={Nucleic Acids Research},
  volume={34},
  pages={W609--W612},
  year={2006}
}

@article{nei1986simple,
  title={Simple methods for estimating the numbers of synonymous and nonsynonymous nucleotide substitutions},
  author={Nei, Masatoshi and Gojobori, Takashi},
  journal={Molecular Biology and Evolution},
  volume={3},
  number={5},
  pages={418--426},
  year={1986}
}

@article{jumper2021highly,
  title={Highly accurate protein structure prediction with AlphaFold},
  author={Jumper, John and others},
  journal={Nature},
  volume={596},
  number={7873},
  pages={583--589},
  year={2021}
}

@article{rego2015py3dmol,
  title={3Dmol.js: molecular visualization with WebGL},
  author={Rego, Nicholas and Koes, David},
  journal={Bioinformatics},
  volume={31},
  number={8},
  pages={1322--1324},
  year={2015},
  doi={10.1093/bioinformatics/btu829}
}

@article{hunter2007matplotlib,
  title={Matplotlib: A 2D graphics environment},
  author={Hunter, John D},
  journal={Computing in Science \& Engineering},
  volume={9},
  number={3},
  pages={90--95},
  year={2007}
}

@article{grantham1974amino,
  title={Amino acid difference formula to help explain protein evolution},
  author={Grantham, Richard},
  journal={Science},
  volume={185},
  number={4154},
  pages={862--864},
  year={1974}
}

@article{harris2020array,
  title={Array programming with NumPy},
  author={Harris, Charles R and others},
  journal={Nature},
  volume={585},
  number={7825},
  pages={357--362},
  year={2020}
}

% NOTA: mejor como inproceedings + DOI oficial de SciPy Proceedings.
@inproceedings{mckinney2010data,
  title={Data Structures for Statistical Computing in Python},
  author={McKinney, Wes},
  booktitle={Proceedings of the 9th Python in Science Conference (SciPy 2010)},
  pages={56--61},
  year={2010},
  doi={10.25080/Majora-92bf1922-00a}
}
