## Supplementary figures and images for "Divergent disulfide bond architecture defines two IgY subclasses in snakes"

### dNdS_sliding_window.png

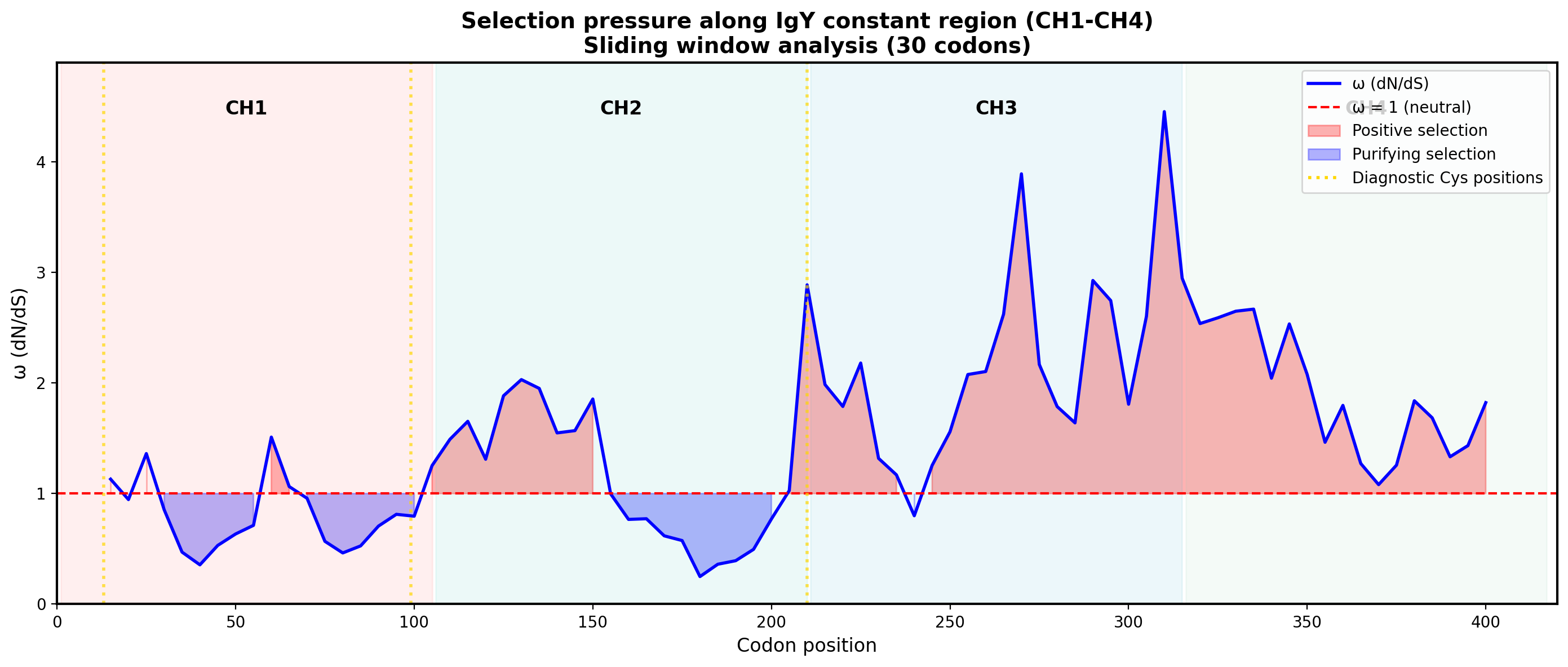

### Figure_IgY_disulfide_scheme_publication.pdf

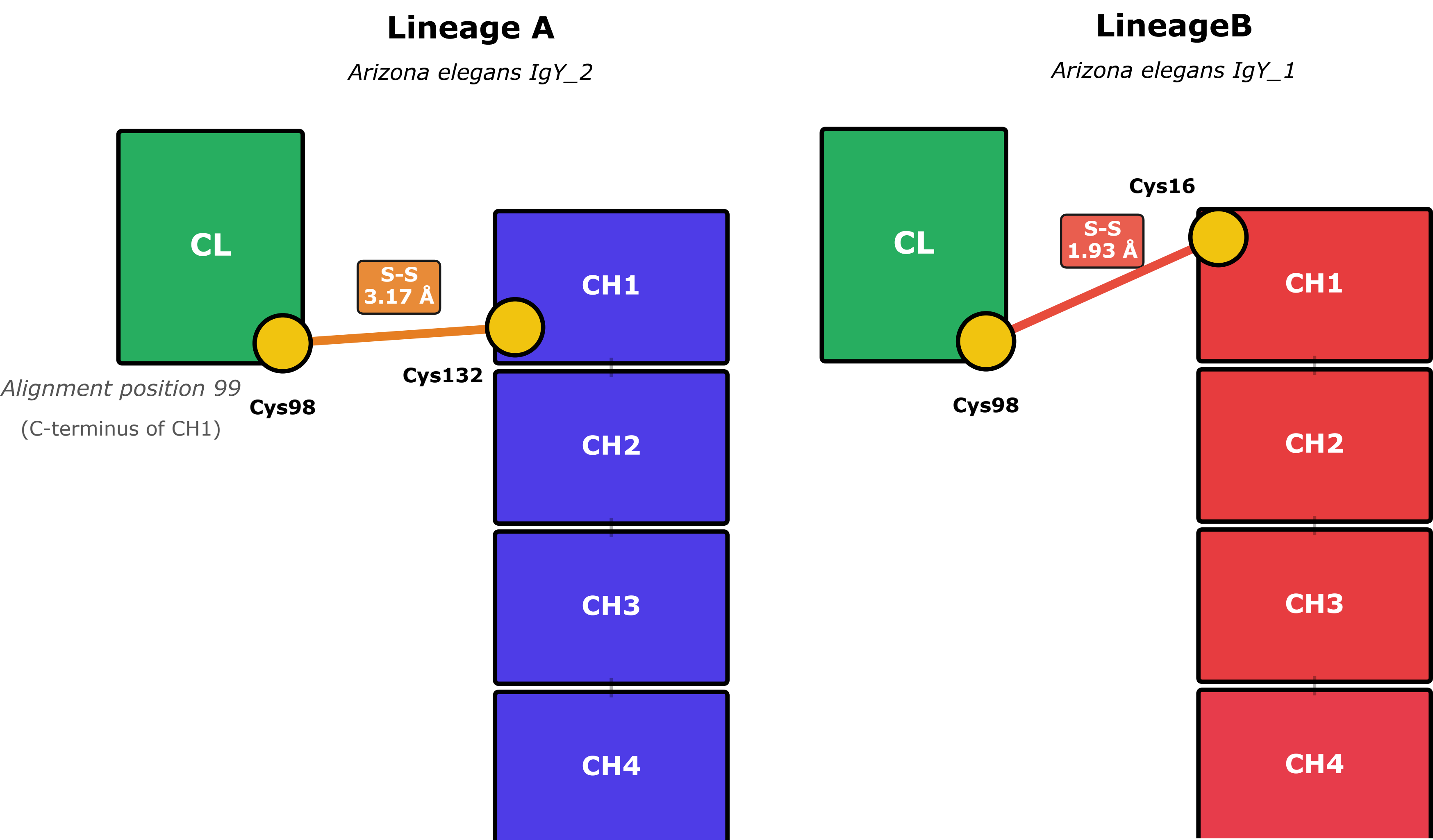

### IgY1.png

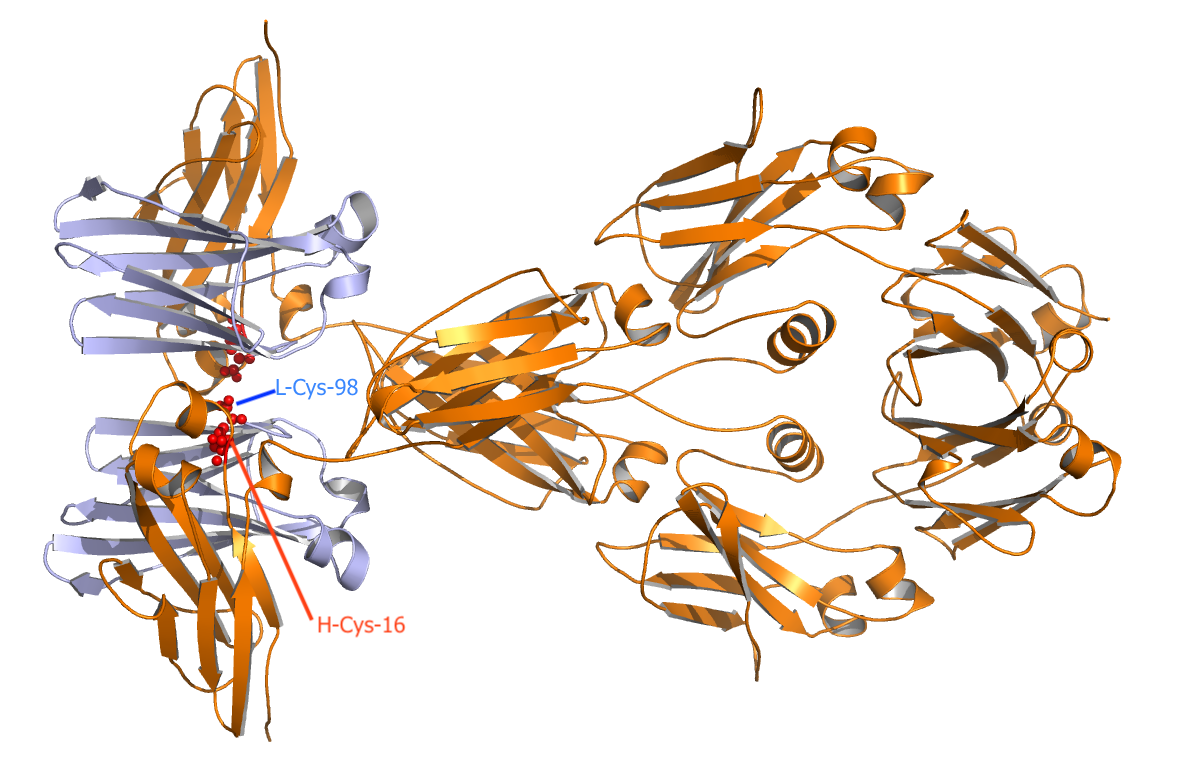

### IgY2.png

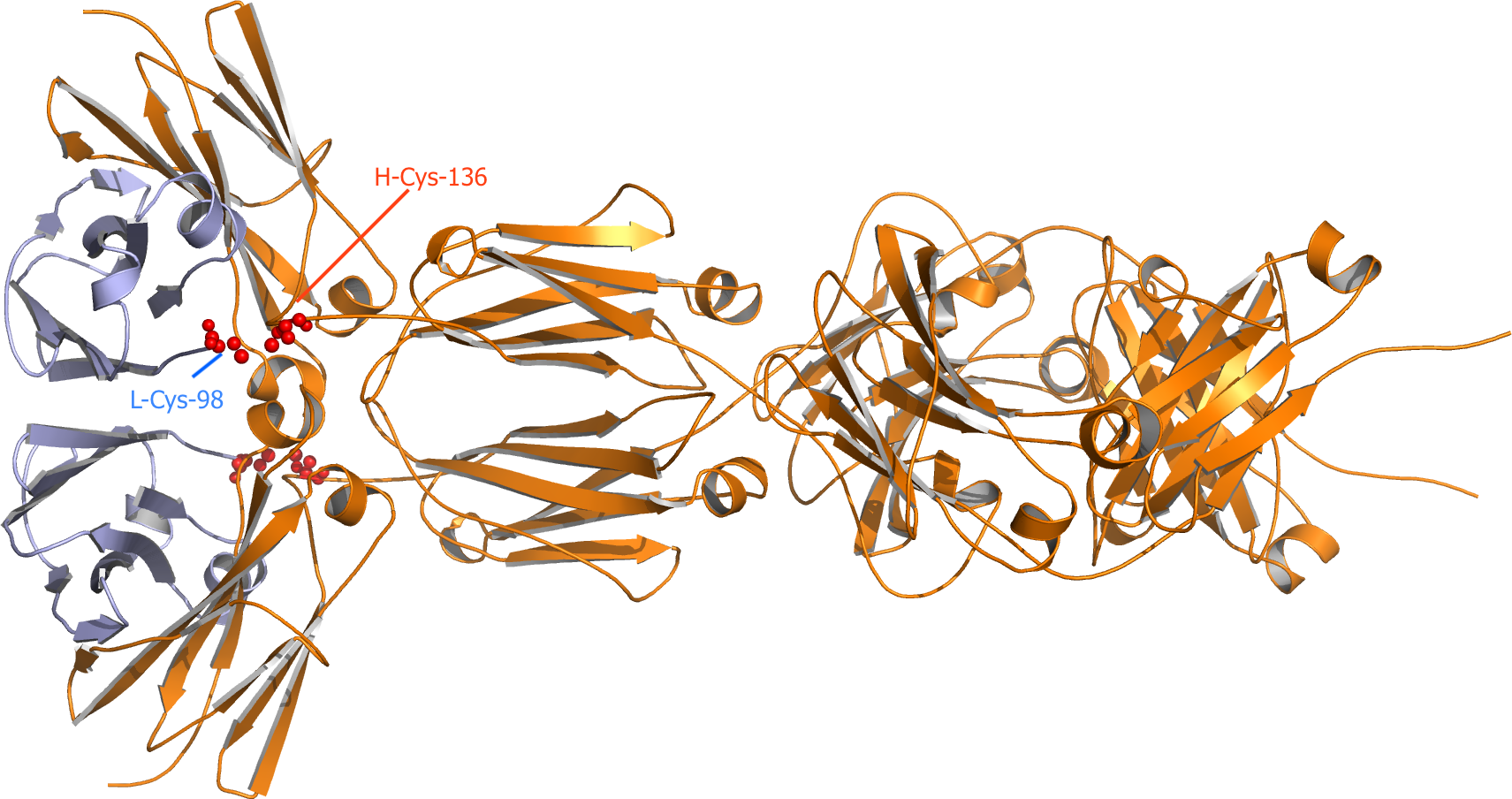

### IgY-with-LC.png

#### IgY1 vs IgY2 – disulfide architecture (HC and LC constant region) inferred from models

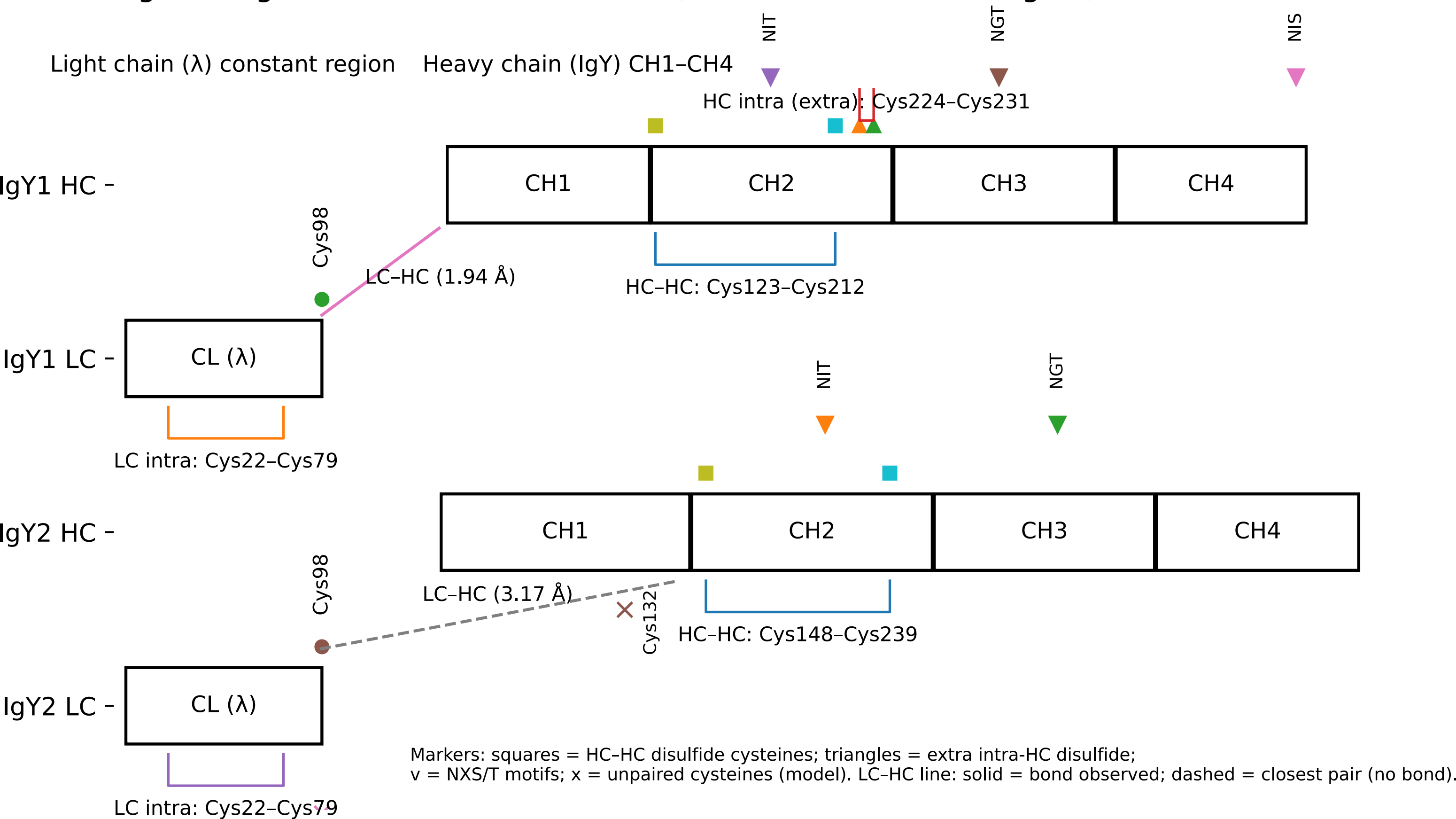

### IgY_all_aa.aln.occ80.fasta.treefile.pdf

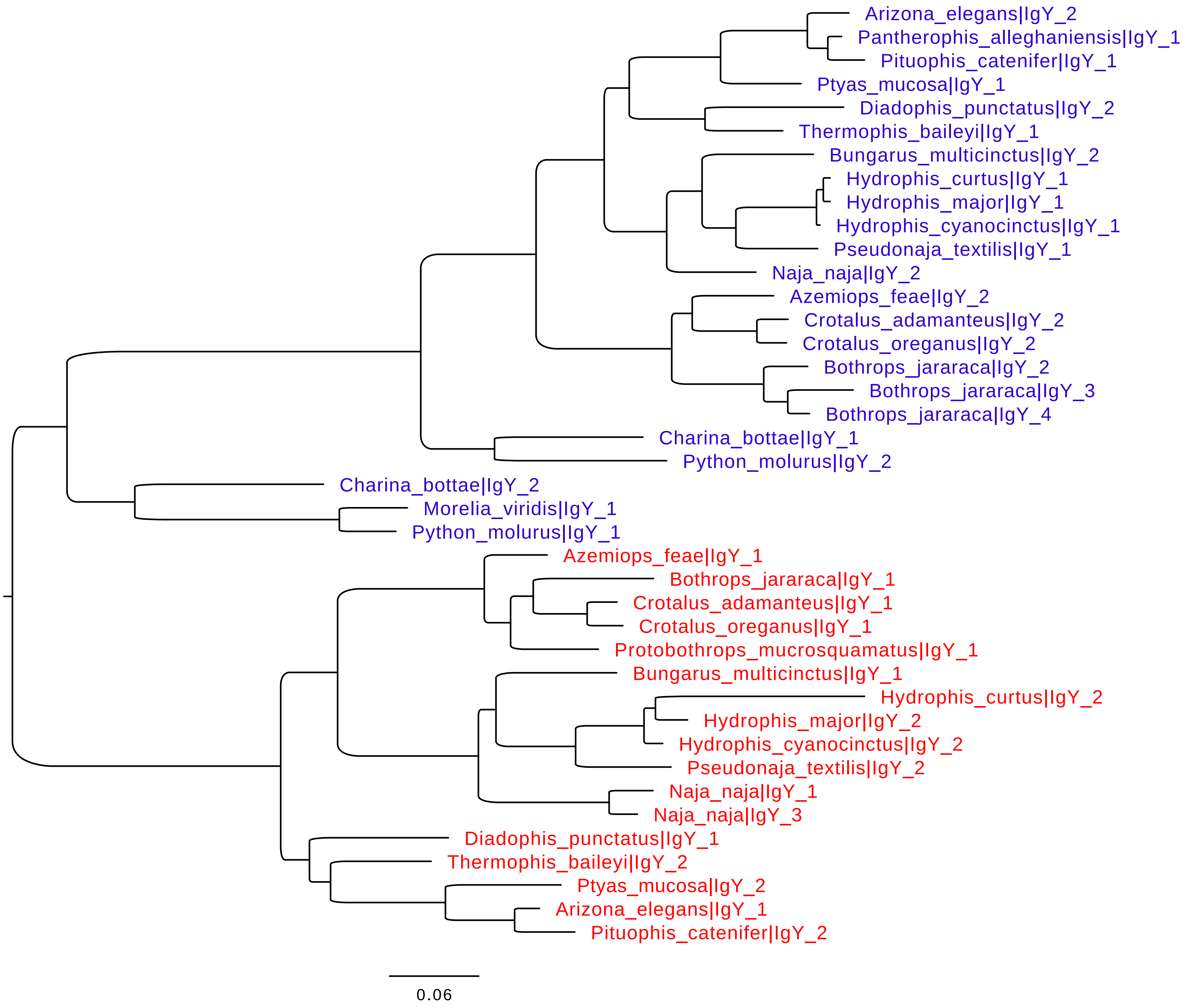
